# Comparative genomics of *Clostridium* species associated with vacuum-packed meat spoilage

**DOI:** 10.1101/2020.09.03.282111

**Authors:** Nikola Palevich, Faith P. Palevich, Paul H. Maclean, Eric Altermann, Amanda Gardner, Sara Burgess, John Mills, Gale Brightwell

**Affiliations:** AgResearch Limited, Grasslands Research Centre, Palmerston North, New Zealand; Riddet Institute, Massey University, Palmerston North, New Zealand; Molecular Epidemiology and Veterinary Public Health Laboratory (mEpiLab), Infectious Disease Research Centre, School of Veterinary Science, Massey University, Palmerston North, New Zealand

**Keywords:** Genomics, Comparative genomics, *Clostridia*, *Clostridium estertheticum*, *Clostridium tagluense*, Vacuum-packageded meat, Spoilage flora, Carbohydrate metabolism

## Abstract

Bacterial species belonging to the *Clostridium* genera have been recognized as causative agents of blown pack spoilage (BPS) in vacuum packed meat products. Whole-genome sequencing of six New Zealand psychrotolerant *Clostridium* isolates derived from three meat production animal types and their environments was performed to examine their roles in BPS. Comparative genome analyses have provided insight into the genomic diversity and physiology of these bacteria and divides *Clostridia* into two separate species clusters. BPS-associated *Clostridia* encode a large and diverse spectrum of degradative carbohydrate-active enzymes (CAZymes). In total, 516 glycoside hydrolases (GHs), 93 carbohydrate esterases (CEs), 21 polysaccharide lyases (PLs), 434 glycosyl transferases (GTs) and 211 carbohydrate-binding protein modules (CBM) with predicted activities involved in the breakdown and transport of carbohydrates were identified. *Clostridia* genomes have different patterns of CAZyme families and vary greatly in the number of genes within each CAZy category, suggesting some level of functional redundancy. These results suggest that BPS-associated *Clostridia* occupy similar environmental niches but apply different carbohydrate metabolism strategies to be able to co-exist and cause meat spoilage.

## 1.0. Introduction

Despite stringent control measures for vacuum-packaging and regulation of storage temperatures for chilled fresh meat products destined for overseas markets, premature blown pack spoilage (BPS) of vacuum packaged meat can still occur. Spore-forming bacteria, namely psychrophilic and psychrotrophic *Clostridium* species (Mills et al., 2014), are common contaminants of food and the environment, and as such represent a major source of food poisoning and food spoilage. The global economic losses attributed to product spoilage and market access issues are significant and lead to reduced consumer confidence.

Numerous bacterial species belonging to the *Clostridium* genera have been associated as causative agents of blown pack spoilage (BPS) in vacuum packed meat products, including: *C. algidicarnis* (Lawson et al., 1994), *C. algidixylanolyticum* (Broda et al., 2000a), *C. gasigenes* (Broda et al., 2000b), *C. bowmanii* (Spring et al., 2003), *C. frigidicarnis* (Broda et al., 1999), *C. estertheticum* (Collins et al., 1992), *C. estertheticum* subspecies *laramiense* (Kalchayanand et al., 1993), *C. frigoris* (Spring et al., 2003) and *C. tagluense* (Suetin et al., 2009). The above-mentioned *Clostridium* species associated with BPS are typically characterized as Gram-positive, slow-growing, spore-forming, psychotrophic anaerobes. Despite the importance of these spoilage microorganisms, studies are challenged by a lack of differential media or straightforward discriminatory methods for the specific identification of different spoilage *Clostridium* species. The currently available molecular tools such as, amplified rDNA (Ribosomal DNA) restriction analysis (ARDRA) and 16*S* rRNA gene sequencing (Brightwell and Horváth, 2018), are predominantly suitable for only pre-screening and community analysis of *Clostridium* species that may be associated with meat production animal types (cattle, sheep and deer) and their environments. With the advent of genome sequencing technology there is an opportunity to improve our basic knowledge of these important food-production and spoilage associated bacteria. Recently, several reference genomes have been made available (Palevich et al., 2020b; Palevich et al., 2020c; Palevich et al., 2020d; Palevich et al., 2020e; Palevich et al., 2020f), as well as those of characterized type strains *C. estertheticum* DSM 8809^T^ (Yu et al., 2016), *C. tagluense* A121^T^ (Suetin et al., 2009) and C. *estertheticum* subsp. *laramiense* DSM 14864^T^ (Palevich et al., 2019c). In this study, we report a detailed primary-level pan-genome comparative analysis on our six isolates with the nine closely related and previously characterised meat spoilage-associated *Clostridium* isolates (*C. estertheticum, C. estertheticum*-like, *C. gasigenes, C. algidicarnis, C. frigidicarnis* and *C. tagluense*), to highlight their enzymatic machinery and metabolic capacities.

## 2.0. Materials and Methods

### 2.1. Bacterial cultivation and growth conditions

The methods for isolation and cultivation of the various meat spoilage associated *Clostridium* species described in the study has been previously detailed in Broda et al. (1998) and Broda et al. (2000b). Strains DSM 14864^T^ and DSM 8809^T^ were acquired from the Leibniz Institute DSMZ-German Collection of Microorganisms and Cell Cultures. All cultures were retrieved from storage, grown anaerobically at 10°C in pre-reduced Peptone, Yeast Extract, Glucose, Starch broth (PYGS) (Lund et al., 1990) and culture purity checked by plating.

### 2.2. Preparation of genomic DNA and whole-genome sequencing

Genomic DNA was extracted from freshly grown cells using a modification of the phenol-chloroform procedure (Bouillaut et al., 2011). Specificity of genomic DNA was verified by automated Sanger sequencing of the 16*S* rRNA gene following PCR amplification from genomic DNA. Total DNA amounts were determined using a NanoDrop^®^ ND-1000 (Thermo Scientific Inc.) and a Qubit Fluorometer dsDNA BR Kit (Invitrogen, USA), in accordance with the manufacturer’s instructions. Genomic DNA integrity was verified by agarose gel electrophoresis and using a 2000 BioAnalyzer (Agilent, USA). The genomic DNA was mechanically sheared using a Nebulizer instrument (Invitrogen) to select fragments of approximately 550 bp. A DNA library was prepared using the Illumina TruSeq™ Nano method and sequenced on the Illumina MiSeq platform with the 2× 250 bp paired-end (PE) reagent kit v2.

### 2.3. Genome assembly and annotation

The quality of the raw reads was checked in FastQC v0.11.5 (https://www.bioinformatics.babraham.ac.uk/projects/fastqc/), the reads were trimmed with Trimmomatic v0.39 (http://www.usadellab.org/cms/?page=trimmomatic). A *de novo* assembly was performed using the A5-miseq pipeline v20169825 with standard parameters (Coil et al., 2014). Initial genome annotation was performed using GAMOLA2 (Altermann et al., 2017) for in preparation for submission to the National Center for Biotechnology Information (NCBI). The software packages Diamond v0.9.21.122 (Buchfink et al., 2015) and InterProScan v5.36-75.0 (Jones et al., 2014) were used to search the NCBI “nr” (non-redundant) database with the resulting protein set imported into BLAST2GO as implemented in the OmicsBox software package v1.1.164 (Conesa et al., 2005), where gene ontology terms and draft annotations were assigned to each protein.

In addition, genomes were annotated by the to the U.S. Department of Energy (DOE) Joint Genome Institute (JGI) Integrated Microbial Genomes (IMG) genome annotation pipeline, via direct submission to the IMG system (Mavromatis et al., 2009). Briefly, protein-coding genes (coding sequence [CDSs]) were identified using the Prodigal v2.6.3 program (102), followed by a round of automated and manual curation using the MGAP v5.0.12 pipeline (Huntemann et al., 2015). Functional annotation and additional analyses were performed within the Integrated Microbial Genomes Expert Review (IMG-ER) platform (Markowitz et al., 2009). All bioinformatics analyses were performed using default settings and parameters.

The genome sequences and associated data for all six *Clostridium* reported in this study were deposited in NCBI under the BioProject accession number PRJNA574489. In addition, the data sets supporting the conclusions of this article are available through the IMG portal (https://img.jgi.doe.gov/).

### 2.4. Comparative analysis of the genome data sets

#### 2.4.1. Average nucleotide identity (ANI) computation

ANI was used as a measure of genetic relatedness based on the gene content between the 15 *Clostridium* genomes. ANI is a measure of nucleotide-level genomic similarity and was carried out using a BLAST approach (ANIm) using the default parameters in the JSpecies software package v. 3.4.8 between each pair of genomes (Richter et al., 2016). To compare the ANIb values, a heat-map was generated using the heatmap.2 function in the gplots library of the statistics software package R (v. 3.5.2). In order to identify species ANI for the 15 *Clostridium* genomes that determine whether the genomes in a pair belong to the same species, only the subset of high-quality genome pairs were utilized and a ANI cutoff of ≥96% was used to define species.

#### 2.4.2. Functional genome distribution (FGD) analysis

FGD is a tool for comparative microbial genomics analysis and interpretation of the genetic diversity of bacteria (Altermann, 2012). FGD investigates the overall similarity levels between microbial genomes, based on the amino acid sequences of their predicted ORFeomes, which correspond to the coding sequences (CDSs) of the genes (open reading frames [ORFs]) in a genome, and ultimately defines the degree of similarity of the genomes. All 14 *Clostridium* genomes were downloaded in FASTA format from the IMG genome database (111), concatenated using a universal spacer-stop-spacer sequence, and automatically annotated using the GAMOLA2 software package (Altermann et al., 2017). The predicted ORFeomes of all genomes were subjected to an FGD analysis, and the resulting distance matrix was imported into MEGA6 (Tamura et al., 2013). The functional genome distribution was visualized using the unweighted pair group method with arithmetic mean (UPGMA) method (Jones et al., 1992). The procedure outlined here and for manual curation of the genome annotations has been detailed by Palevich et al. (Palevich et al., 2019a; Palevich et al., 2017; Palevich et al., 2019b; Palevich et al., 2020a).

#### 2.4.3. Determination of the core and pan-genomes

The genes representative of the *Clostridium* core and pan-genomes were determined by performing a BLAST-based analysis using OrthoVenn v.2 (Wang et al., 2015) with default parameters, to compare the orthologous gene clusters within the *Clostridium* genomes. Briefly, if two proteins within a genome met the designated cut-off, they were clustered into one protein family. Protein families were extended via single-linkage clustering. If a protein family included proteins from all genomes in the comparison, the family was designated a core protein family. Subset genes, such as species group shared and unique subsets of genes within individual genomes, were identified by clustering the results from the core and pan-genome calculations.

#### 2.4.4. CAZyme annotation

The putative proteomes of the 14 *Clostridium* data sets were subjected to automated annotation and assignment to CAZymes using the dbCAN resource CAZy family-specific hidden Markov models (HMMs) (104). An E value of <1e^-3^ for CAZymes based on family-specific HMMs was used as the cutoff for alignments shorter than 80 amino acids, while an E value of <1e^-5^ was used for alignments longer than 80 amino acids. These cut-off settings enabled short but significant CBM matches to be maintained. All dbCAN hits were clustered at a 100% sequence identity threshold using the CD-HIT Illumina algorithm to remove duplicates (Marchler-Bauer et al., 2012). All descriptions and classifications were compiled from CAZy (Cantarel et al., 2009), and the modular architectures of CAZymes and predicted proteins with multimodular CAZyme organizations in the genome data sets were determined by searching each query protein against the Pfam and Protein Data Bank (PDB) databases (Finn et al., 2013; Rose et al., 2013).

## 3.0. Results

### 3.1. Comparative genomics

NZ strain representatives; *C. tagluense* FP1 and *C. tagluense* FP2, *C. estertheticum* FP3 and FP4, *Clostridium* sp. M14, and type strain *C. estertheticum* subsp. *laramiense* DSM 14864^T^ were selected for genome sequencing to examine their roles in BPS. The *de novo* assemblies of the six *Clostridium* genomes were all determined using Illumina MiSeq technology to generate on average 129 scaffolds with 172× coverage. The N50 values ranged between 43,169 bp (*C. estertheticum* FP4) to 757,921 bp (*Clostridium* sp. M14) with the largest scaffold length being 1,669,648 bp (*Clostridium* sp. M14) and smallest of 133,780 bp (*C. estertheticum* FP4) in size. The draft genome sequences were composed of on average 4,924,744 bp and %G+C content of 30.4% (Table 1). The presented *Clostridium* pan-genome consists of a total of 30,427 putative protein-coding genes (CDS) were predicted along with 21,420 genes with putative functions and an average of approximately 30% of CDS with unknown function predictions. Although the *Clostridium* pan-genome consists of only draft genomes, the presence of extrachromosomal elements (i.e. plasmids, megaplasmids and/or chromids) have been identified with the presence of plasmid replication initiation genes (*rep*) in all but the *C. estertheticum* FP3 genome. Furthermore, *rep* genes were identified in three or more gene clusters and on separate scaffolds in *C. estertheticum* subsp. *laramiense* DSM 14864^T^, *C. tagluense* FP1 and *C. tagluense* FP2, indicating the presence of multiple plasmids.

**Table 1.** Comparison of assembly and annotation statistics for the currently available BPS-associated *Clostridia* genomes.

|  | *C. tagluense* FP1 |  | *C. tagluense* FP2 |  | *C. estertheticum* FP3 |  | *C. estertheticum* FP4 |  | *Clostridium* sp. M14 |  | *C. estertheticum* subsp.*laramiense* DSM 14864<sup>T</sup> |  | *C. estertheticum* DSM 8809<sup>T</sup> |  |
| --- | --- | --- | --- | --- | --- | --- | --- | --- | --- | --- | --- | --- | --- | --- |
|  | Value | % total<sup>a</sup> | Value | % total<sup>a</sup> | Value | % total<sup>a</sup> | Value | % total<sup>a</sup> | Value | % total<sup>a</sup> | Value | % total<sup>a</sup> | Value | % total<sup>a</sup> |
| Genome Project Information |  |  |  |  |  |  |  |  |  |  |  |  |  |  |
| Status | Draft |  | Draft |  | Draft |  | Draft |  | Draft |  | Draft |  | Complete |  |
| Isolation source | Lamb |  | Venison |  | Lamb |  | Lamb |  | Venison |  | Beef |  | Beef |  |
| BioSample ID | SAMN14128649 |  | SAMN14128650 |  | SAMN14128651 |  | SAMN14128652 |  | SAMN14128653 |  | SAMN12859418 |  | SAMN04958870 |  |
| BioProject ID |  |  |  |  |  |  | PRJNA574489 |  |  |  |  |  | PRJNA320887 |  |
| Assembly method | A5-miseq v. 20169825 |  | A5-miseq v. 20169825 |  | A5-miseq v. 20169825 |  | A5-miseq v. 20169825 |  | A5-miseq v. 20169825 |  | A5-miseq v. 20169825 |  | HGAP2 v. 26/11/2015 |  |
| Genome coverage | 145x |  | 115x |  | 139x |  | 242x |  | 184x |  | 202x |  | 522x |  |
| Sequencing technology |  |  |  |  |  |  | Illumina MiSeq |  |  |  |  |  | PacBio RS II |  |
| Annotation Method | IMG Annotation Pipeline v.5.0.15 |  |  |  |  |  |  |  |  |  |  |  |  |  |
| Genome Statistics |  |  |  |  |  |  |  |  |  |  |  |  |  |  |
| Genome size (bp) | 5,379,343 |  | 5,549,561 |  | 5,555,543 |  | 4,088,187 |  | 3,981,244 |  | 4,994,588 |  | 4,785,613 |  |
| DNA coding (bp) | 4,498,050 | 83.6 | 4,630,094 | 83.4 | 4,657,543 | 83.8 | 3,422,892 | 83.7 | 3,326,305 | 83.6 | 4,260,218 | 85.3 | 4,123,313 | 86.2 |
| DNA G+C (bp) | 1,671,864 | 31.1 | 1,715,103 | 30.9 | 1,750,446 | 31.5 | 1,278,166 | 31.3 | 1,081,186 | 27.2 | 1,525,815 | 30.6 | 1,478,579 | 30.9 |
| DNA replicons/scaffolds | 209 |  | 110 |  | 85 |  | 251 |  | 35 |  | 84 |  | 2 |  |
| Genome Annotations |  |  |  |  |  |  |  |  |  |  |  |  |  |  |
| Total genes | 5,705 |  | 5,661 |  | 5,685 |  | 4,302 |  | 3,958 |  | 5,116 |  | 4,656 |  |
| Protein coding genes (CDS) | 5,373 | 94.2 | 5,376 | 95.0 | 5,434 | 95.6 | 4,027 | 93.6 | 3,768 | 95.2 | 4,889 | 95.6 | 4,498 | 96.6 |
| RNA genes | 197 | 3.5 | 185 | 3.3 | 168 | 3.0 | 153 | 3.6 | 126 | 3.2 | 149 | 2.9 | 158 | 3.4 |
| rRNA genes | 91 | 1.6 | 80 | 1.4 | 59 | 1.0 | 55 | 1.3 | 41 | 1.0 | 54 | 1.1 | 48 | 1.0 |
| tRNA genes | 100 | 1.8 | 98 | 1.7 | 104 | 1.8 | 94 | 2.2 | 81 | 2.0 | 91 | 1.8 | 89 | 1.9 |
| CDS genes with function prediction | 3,863 | 67.7 | 3,956 | 69.9 | 4,106 | 72.2 | 3,052 | 70.9 | 2,789 | 70.5 | 3,654 | 71.4 | 3,390 | 72.8 |
| CDS genes with unknown function | 1,510 | 26.5 | 1,420 | 25.1 | 1,328 | 23.4 | 975 | 22.7 | 979 | 24.7 | 1,235 | 24.1 | 1,108 | 23.8 |
| Genes with enzymes | 986 | 17.3 | 1,030 | 18.2 | 1,102 | 19.4 | 841 | 19.6 | 845 | 21.4 | 1,055 | 20.6 | 1,040 | 22.3 |
| Genes connected to KEGG pathways | 1,042 | 18.3 | 1,116 | 19.7 | 1,192 | 21.0 | 873 | 20.3 | 911 | 23.0 | 1,166 | 22.8 | 1,138 | 24.4 |
| Genes connected to KEGG Orthology (KO) | 2,050 | 35.9 | 2,137 | 37.8 | 2,219 | 39.0 | 1,745 | 40.6 | 1,634 | 41.3 | 2,123 | 41.5 | 2,049 | 44.0 |
| Genes connected to MetaCyc pathways | 866 | 15.2 | 902 | 15.9 | 973 | 17.1 | 748 | 17.4 | 742 | 18.8 | 931 | 18.2 | 921 | 19.8 |
| Genes assigned to COGs | 3,776 | 66.2 | 3,843 | 67.9 | 4,015 | 70.6 | 2,997 | 69.7 | 2,706 | 68.4 | 3,668 | 71.7 | 2,800 | 60.1 |
| Genes with Pfam domains | 3,891 | 68.2 | 4,005 | 70.8 | 4,121 | 72.5 | 3,031 | 70.5 | 2,803 | 70.8 | 3,738 | 73.1 | 3,508 | 75.3 |
| Genes with TIGRfam domains | 1,315 | 23.1 | 1,347 | 23.8 | 1,333 | 23.5 | 1,135 | 26.4 | 1,131 | 28.6 | 1,344 | 26.3 | 1,336 | 28.7 |
| Genes with signal peptides | 142 | 2.5 | 151 | 2.7 | 148 | 2.6 | 86 | 2.0 | 114 | 2.9 | 146 | 2.9 | 197 | 4.2 |
| Genes with transmembrane helices | 1,327 | 23.3 | 1,381 | 24.4 | 1,370 | 24.1 | 1,050 | 24.4 | 910 | 23.0 | 1,331 | 26.0 | 1,232 | 26.5 |
| Horizontally transferred genes | 210 | 3.7 | 252 | 4.5 | 364 | 6.4 | 201 | 4.7 | 61 | 1.5 | 35 | 0.7 | 209 | 4.5 |
| CRISPR count | 4 |  | 2 |  | - |  | 2 |  | 6 |  | - |  | - |  |
| Gene Cassettes and Clusters |  |  |  |  |  |  |  |  |  |  |  |  |  |  |
| Chromosomal Cassette count | 775 |  | 728 |  | 759 |  | 629 |  | 568 |  | 606 |  | 505 |  |
| Chromosomal Cassette genes | 5,411 | 94.9 | 5,376 | 95.0 | 5,445 | 95.8 | 4,060 | 94.4 | 3,771 | 95.3 | 4,985 | 97.4 | 4,564 | 98.0 |
| COG clusters | 1,787 | 31.3 | 1,776 | 31.4 | 1,826 | 32.1 | 1,588 | 36.9 | 1,592 | 40.2 | 1,791 | 35.0 | 1,518 | 32.6 |
| Pfam clusters | 2,100 | 36.8 | 2,105 | 37.2 | 2,135 | 37.6 | 1,866 | 43.4 | 1,872 | 47.3 | 2,103 | 41.1 | 2,069 | 44.4 |
| TIGRfam clusters | 964 | 16.9 | 972 | 17.2 | 972 | 17.1 | 897 | 20.9 | 915 | 23.1 | 1,008 | 19.7 | 1,078 | 23.2 |
| Reference | Palevich et al. (2020c) |  | Palevich et al. (2020e) |  | Palevich et al. (2020a) |  | Palevich et al. (2020b) |  | Palevich et al. (2020d) |  | Palevich et al. (2019) |  | Yu et al. (2016) |  |

<sup>a</sup>The total is based on either the size of the genome in base pairs or the total number of genes or CDS in the annotated genome.

Comparative genome analyses were carried out on the six *Clostridium* strains isolated from our lab, along with an additional eight *Clostridium* strains representative of species associated with spoilage of meat (Table 1). Functional genome distribution (FGD) and average nucleotide identity (ANI) were used to investigate the phylogenomic relationships. To examine the taxonomic classification of these *Clostridium* spp., the ANIb values were calculated between each pair of genomes and visualized using a heatmap (Fig. 1A). Those ANIb values greater than 96% were enclosed by a red box, grouping them within the same taxon. The meat spoilage strains currently designated as *C. estertheticum* FP3, *C. estertheticum* FP4 and *Clostridium* sp. M14, all had ANIb values of less than 96% against all of the type strains, suggesting they are novel taxa.

**Figure 1.**
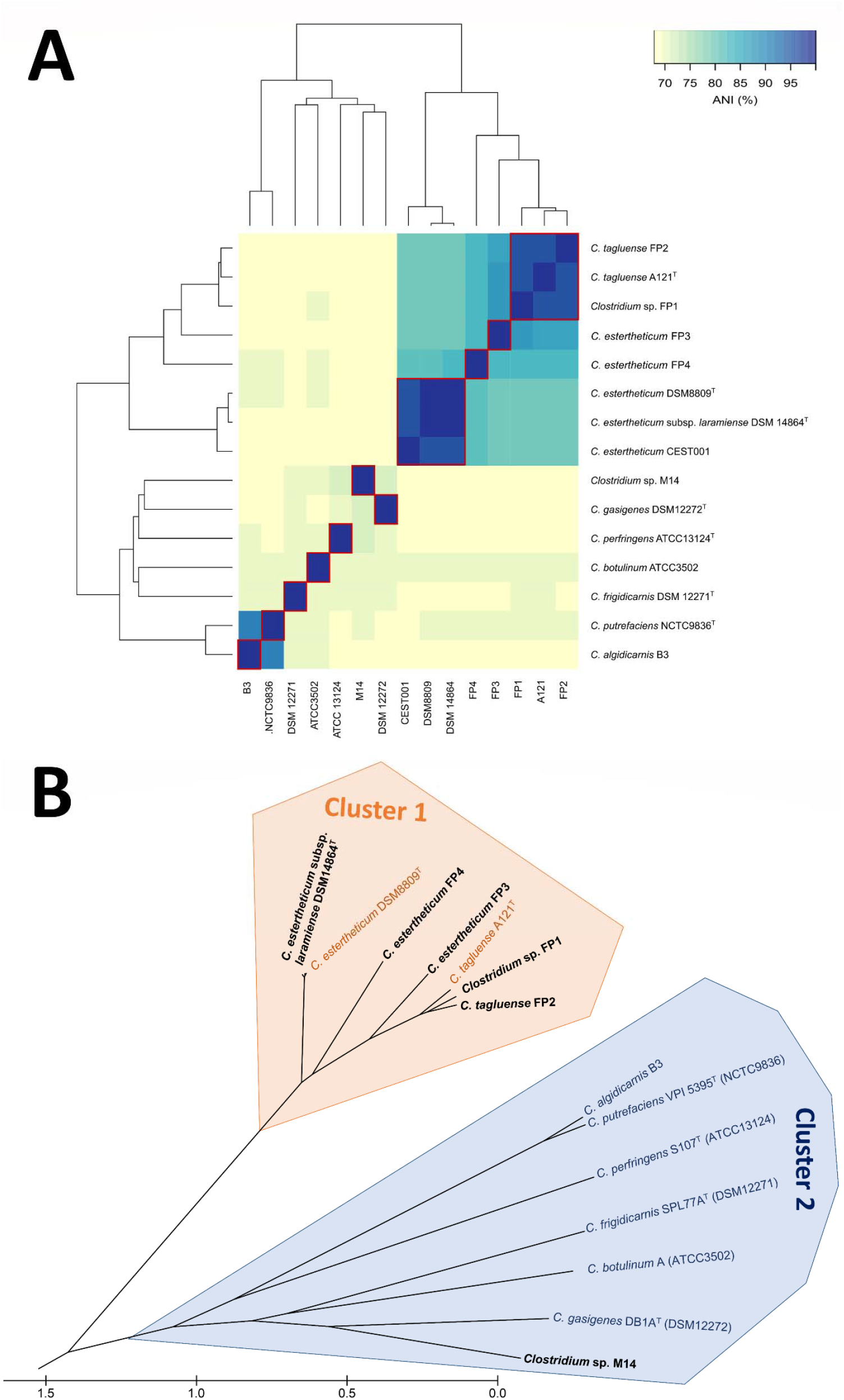
(A) Heatmap based on average nucleotide identity (ANIb) between genomes of BPS-associated *Clostridia* (C.) strains. Dendrogram at the top and on the left are based on reciprocal pairwise comparison clustering calculated using Jspecies (Richter et al., 2016) and visualized using the heatmap.2 function in the R package ggplots2. Those ANIb values greater than 96%, grouping the *Clostridia* strains within the same taxon, were enclosed by a red box. (B) Functional genome distribution (FGD) analysis of BPS-associated *Clostridia*. The predicted ORFeomes of all 15 genomes were subjected to an FGD analysis (Altermann, 2012), and the resulting distance matrix was imported into MEGA6 (Tamura et al., 2013). The functional distribution was visualized using the UPGMA method (Jones et al., 1992). The tree is drawn to scale, with the branch lengths being in the same units as those of the functional distances used to infer the distribution tree. The bar represents the number of nucleotide substitutions per site.

The findings of the FGD analysis grouped all *Clostridium* species into two clusters, that placed *C. estertheticum* and *C. tagluense* strains together and clustered separately from all other *Clostridium* strains (Fig. 1B). Cluster 1 contained the sequences of the type strains of *C. estertheticum* (DSM 8809^T^), *C. estertheticum* subsp. *laramiense* (DSM 14864^T^) and *C. tagluense* (A121^T^) and four other *Clostridium* strains. Within Cluster 1, the currently designated *C. estertheticum* FP3 and especially FP4 clustered separately from the *C. estertheticum* and *C. tagluense* strains that may well represent new *Clostridium* species. Cluster 2 consisted of the sequences of seven *Clostridium* strains containing numerous type strains that represent other *Clostridia* associated with meat production but not BPS, in which *Clostridium* sp. M14 was found to be closely related to the *C. gasigenes* (DB1A^T^) type strain. Clusters 1 and 2 were well supported by bootstrap analyses, while Cluster 2 was more diverse, suggesting that the *Clostridium* strains can be divided into one relatively cohesive cluster (Cluster 1), while the larger Cluster 2 is a continuum of related species (Fig. 1).

The core, variable, and unique gene families present in the Cluster 1 *Clostridium* genomes were determined using BLAST analyses. Overall, 751 gene clusters, 545 orthologous clusters (at least contains two species) and a total of 206 single-copy orthologous gene families were found (Fig. 2), of which 292 represented the gene families shared among all five genomes, also referred to as the core genome set. The core genome set consisted mainly of genes encoding housekeeping, carbohydrate metabolism, and transport functions. The *C. estertheticum* subsp. *laramiense* (DSM 14864^T^) and *C. estertheticum* FP3 genomes had the highest number of unique genes (n=26 and 29), with predicted functions including sequence-specific DNA binding (GO:0043565;) and DNA restriction-modification system (GO:0009307).

**Figure 2.**
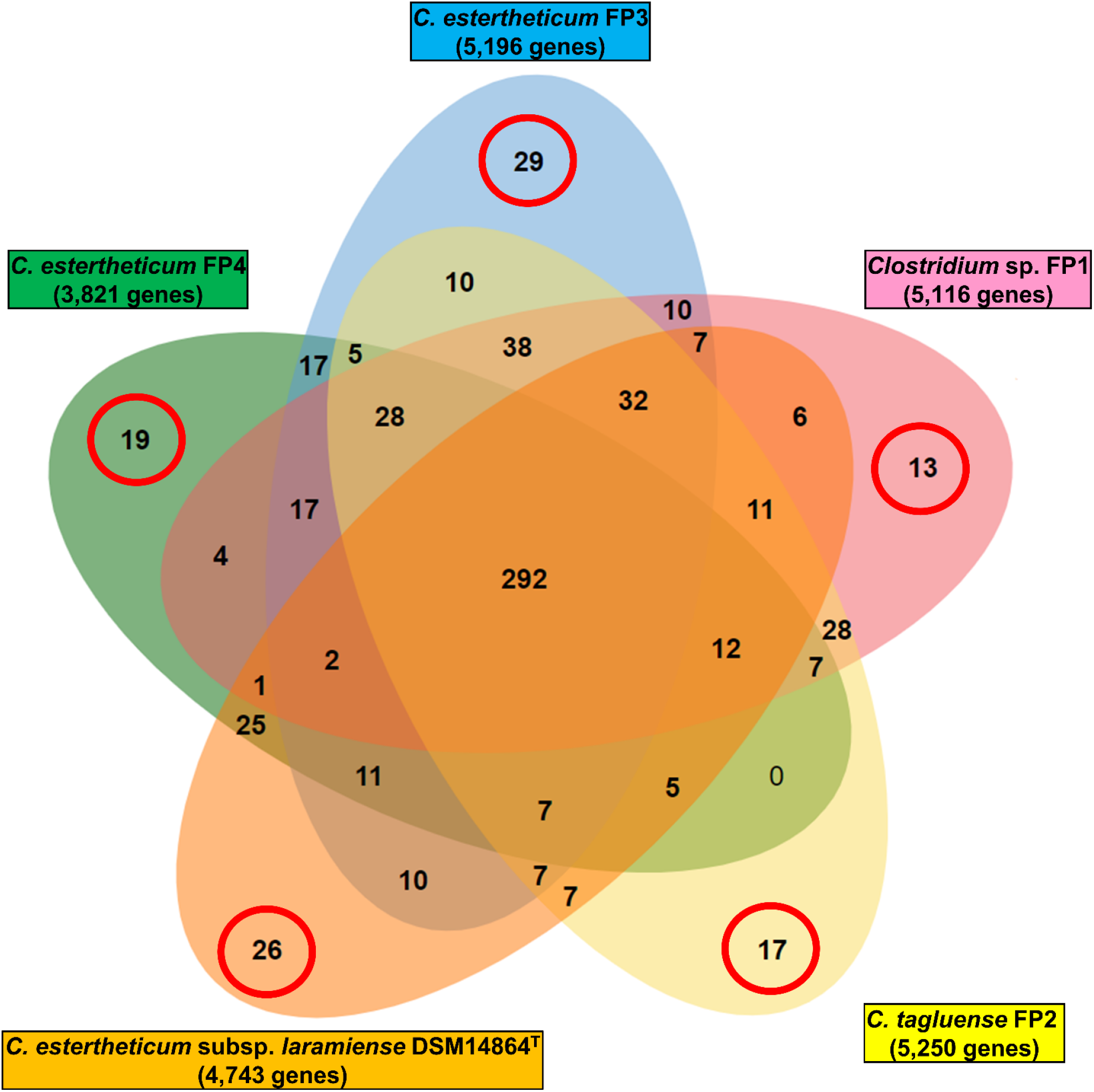
Venn diagram showing the distribution of unique, group-specific, and core gene families among the cluster 1 *Clostridia* genomes. All *Clostridia* scaffolds with at least a single one-to-one ortholog shared among the genomes were compared using OrthoVenn v2 (Wang et al., 2015). The core genome is shown in the center circle. Each coloured intersect segment represents the number of gene families shared among the respective overlapping genomes, and the outermost numbers circled in red represent unique gene families for individual genomes.

### 3.2. Carbohydrate-Active enZYme (CAZyme) profiling

CAZyme profiling was analyzed using dbCAN2 (Zhang et al., 2018) and revealed that the *Clostridium* pan-genome is predicted to encode a total of 516 glycoside hydrolases (GHs), 93 carbohydrate esterases (CEs), 21 polysaccharide lyases (PLs), 434 glycosyl transferases (GTs) and 211 carbohydrate-binding protein modules (CBM) families (Fig. 3). Within the *Clostridium* species, the strains generally had similar types of CAZymes, but with large variations in the absolute numbers of genes within each of their categories in the CAZy profiles. Overall, approximately 2% of the *Clostridium* pan-genome (483 CDSs) is predicted to encode either secreted (70) or intracellular (413) proteins dedicated to carbohydrate and even polysaccharide degradation. Pfam domain analysis of the most abundant GH (GH18, GH3, GH73, and GH13) and CE4 families showed that most did not contain signal peptide sequences and hence predicted to be located intracellularly. Interestingly, the enzymatic profiles of DSM 14864^T^ and the well-characterized *C. estertheticum* DSM 8809^T^ (ATCC 51377^T^) are almost identical, the pair was also atypical of those of their closest Cluster 1 relatives and were separated by CAZyme analysis. Also, similarities were observed among the *C. estertheticum* FP3 and FP4, also *C. tagluense* FP2 and *Clostridium* sp. FP1 pairs of strains (Fig. 3). In addition, *Clostridium* sp. M14 had the most unusual CAZy profile that appeared to be similar to that of the *C. estertheticum* strains, but with a particularly large number of CEs.

**Figure 3.**
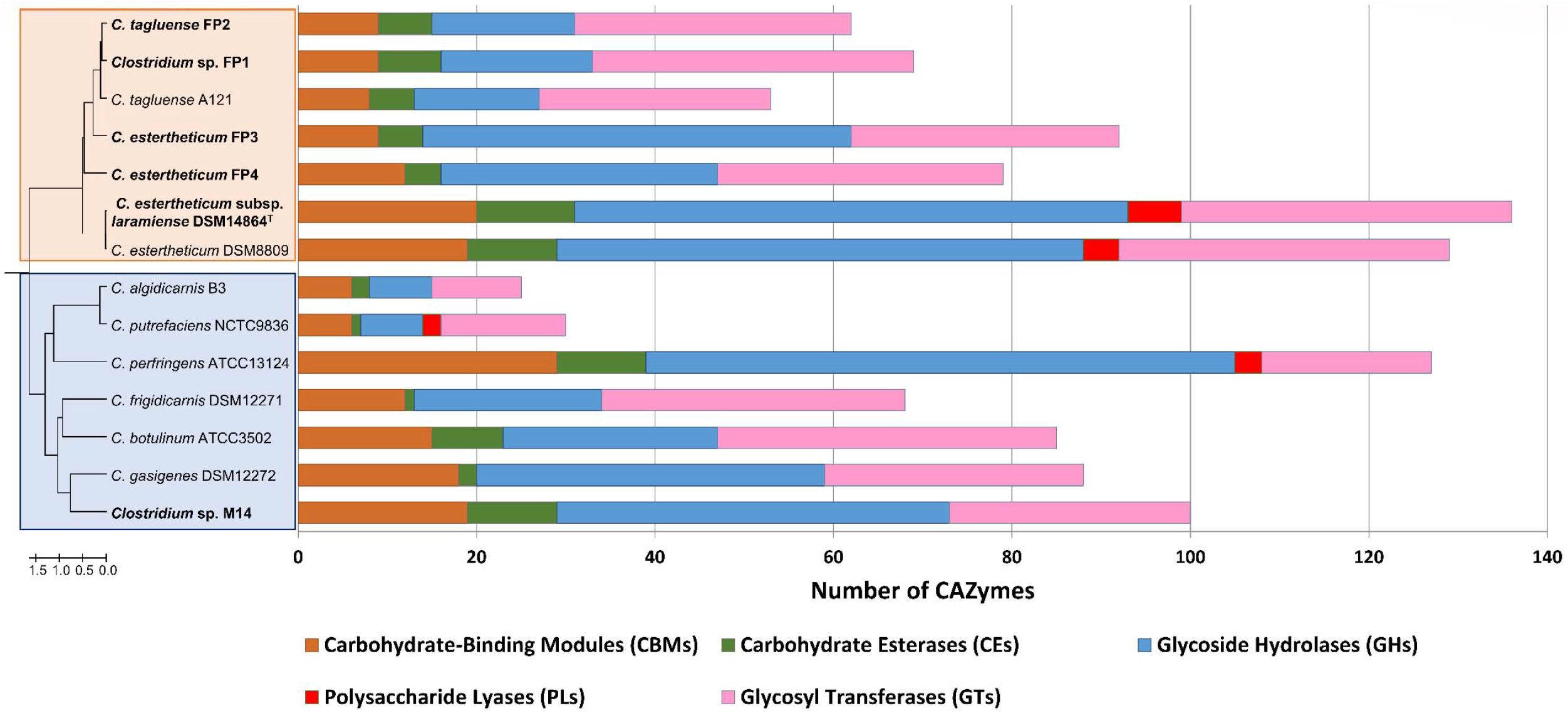
Carbohydrate-Active Enzyme (CAZy) profiles of annotated *Clostridia* genomes. Analysis of the CAZy profiles were annotated using the dbCAN2 resource CAZy family-specific hidden Markov models (HMMs) (Zhang et al., 2018). The numbers and types of CAZyme modules or domains are represented as coloured horizontal bars.

### 3.3. Pathway analysis of carbohydrate metabolism

The fermentation pathways in meat spoilage associated *Clostridium* predicted from gene content and metabolic pathway reconstruction are shown in Fig. 4. Overall, all of the genes encoding the enzymes required for fermenting hexoses through to pyruvate *via* an intact Embden-Meyerhof-Parnas (EMP) pathway were identified in the *Clostridium* pan-genome. The complete methylglyoxal shunt pathway for the alternative production of lactate and mediated by the enzymes: fructose-1,6-bisphophate aldolase (*fbp*), methylglyoxal synthase (*mgsA*), glyoxylase (*gloA/B*), S-lactoylglutathione hydrolase, and _D_-lactate dehydrogenase (*ldhD*), converting _D_-fructose-1,6-bisphophate to pyruvate (Fig. 4), was also investigated. Although lactate production as a fermentation end product was not assessed as part of this study, an incomplete set of methylglyoxal shunt pathway genes was only reported for *Clostridium* sp. M14 (lactate racemase). The genes encoding lactate dehydrogenase (*ldh*) have been identified and compared, in which the *ldh* gene encoding _L_-lactate dehydrogenase plays a key role in the production of _L_-lactate from pyruvate was present in all *Clostridium* genomes, but not *ldhD*.

**Figure 4.**
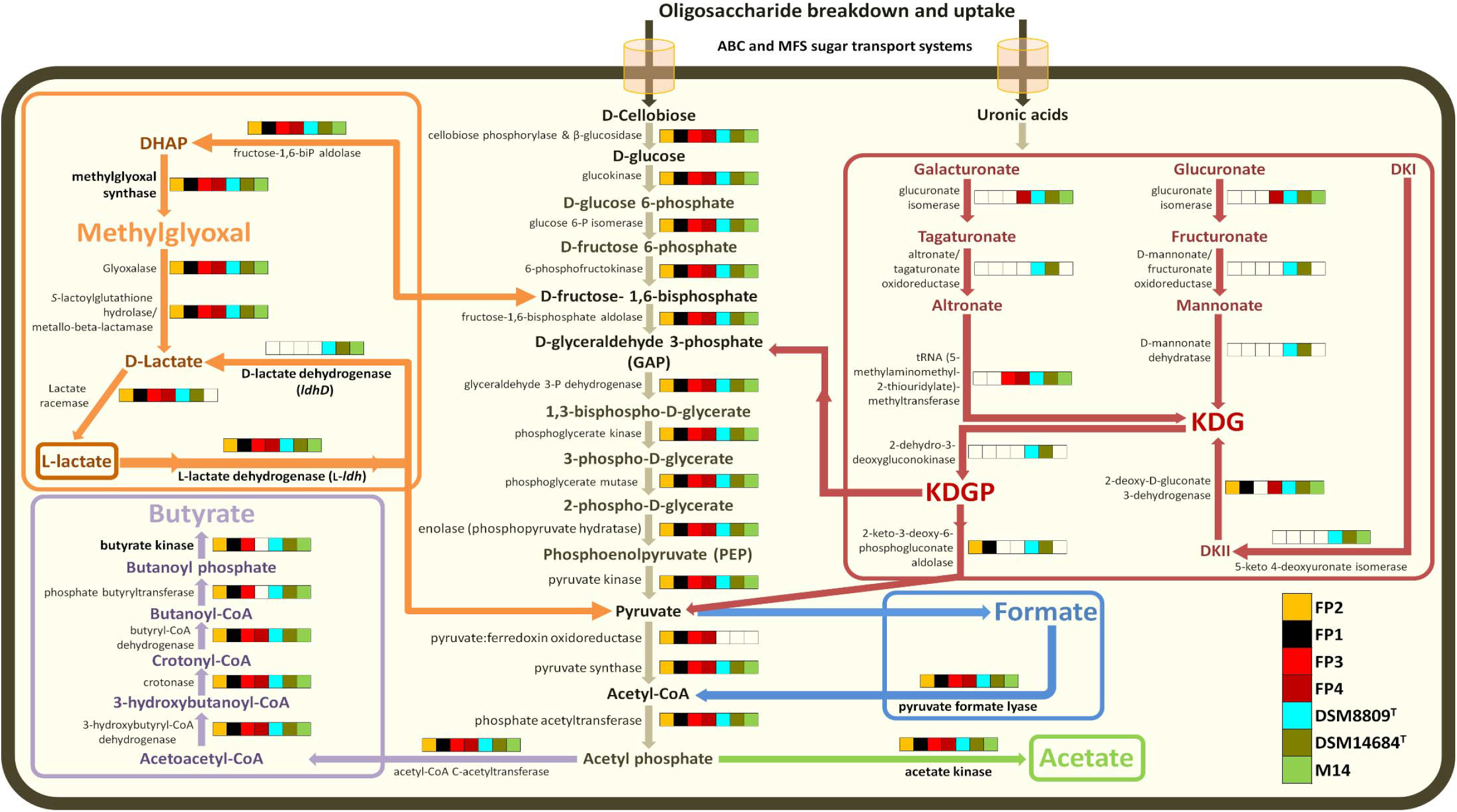
Comparisons of gene presence/absence for enzymes involved in the carbohydrate metabolic pathways in *Clostridia* leading to the formation of butyrate, formate, acetate, and lactate. All metabolic pathways were compiled using information from the MetaCyc (Caspi et al., 2012) and KEGG (Kanehisa and Goto, 2000) databases. The presence or absence of genes encoding particular enzymes within genomes is indicated by full or empty cells, respectively in the panels. The order of genomes in the panels is described in the bottom right corner. Color schemes for the metabolism pathways are as follows: the formation of formate in blue, acetate in green, butyrate in purple, _L_-lactate in red, and _D_-lactate by the proposed methylglyoxal shunt in orange (Cooper, 1984). Abbreviations: DHAP, dihydroxyacetone phosphate; DKI, 5-keto-4-deoxyuronate; DKII, 2,5-diketo3-deoxygluconate; KDG, 2-keto-3-deoxygluconate; KDGP, 2-keto-3-deoxy-gluconate phosphate. Abbreviations for sugar transport systems are as follows: ABC, ATP binding cassette; MFS, major facilitator superfamily.

## 4.0 Discussion

Approximately 200 clostridial species are currently recognized with at best only 10% of these validly characterized due to their phenotypic and metabolic similarities, but also due to the time-consuming and inconsistent cultural differentiation (Broda et al., 2003; Collins et al., 1994; Yutin and Galperin, 2013). Members of the genus *Clostridium* considered to be the major components of the meat spoilage-associated microflora are currently divided into four species, represented by *C. estertheticum, C. gasigenes, C. algidicarnis*, and *C. tagluense*, (Brightwell and Clemens, 2012; Broda et al., 2002; Broda et al., 2009). All of these species belong to the genetically diverse Clostridiaceae family, within the order Clostridiales (Vos et al., 2011). To date, the FDG and ANI analyses reported in this work have provided the highest resolution of the phylogenomic associations for meat spoilage-associated *Clostridium* (Fig. 1), that have further highlighted previously reported inconsistencies between the 16*S* rRNA gene sequence and RFLP (restriction fragment length polymorphism) data (Brightwell and Horváth, 2018). Cluster 1 was phylogenetically cohesive, while the larger Cluster 2 appears to contain a continuum of related organisms.

The pan-genome analysis has revealed that while the Cluster 1 *Clostridium* genomes share about 300 core genes, they also carry unique selections of genes drawn from the species’ accessory genomes. Recently it has been proposed that gene loss and consequently gene gain *via* lateral transfer and gene duplication account for gene loss/gain and may occur at higher rates in organisms on the tips of the phylogenetic tree (Mcinerney et al., 2017). Examples of *Clostridium* strains that may exhibit such genome level plasticity include Cluster 1 strains *C. estertheticum* FP3 and FP4, and Cluster 2 strain *Clostridium* sp. M14. The collective genome cluster complement (3,495 genes) and the core genome (292 genes) of the cluster 1 *Clostridium* strains reflect a large reservoir of genetic diversity within this group (Fig. 2). The strict core genome represents 8.4% of the collective genome and represents the proposed minimum set of genes that allow the survival of *Clostridium* species in the vacuum-packaged meat products. The core genome includes genes encoding protein processing, folding and secretion, cellular processes, energy metabolism and numerous poorly characterized genes (conserved hypothetical proteins, etc.).

The presence of an extrachromosomal element has recently been described for the *C. estertheticum* DSM 8809^T^ type strain reference genome with the presence of a single 23,034 bp plasmid pDSM8009 (Yu et al., 2016). The present study has revealed that extrachromosomal plasmids are common in *Clostridium* species associated with meat spoilage. However, as these findings are based on strictly *in silico* analysis, further experimental validation will be required to confirm these findings such as *via* pulsed-field gel electrophoresis (PFGE) and with additional sequencing to improve the resolution of our genomes (Palevich, 2011; Palevich, 2016; Palevich et al., 2019b). The metabolic burden associated with sustaining the plasmid are made worthwhile for the host as they are likely to play a role as a channel for the horizontal exchange of genomic material and conveying advantageous functions (Jain et al., 2003). The types of essential traits that are transferred by plasmids include those implicated in amino acid, protein, and carbohydrate metabolism, as well as genes encoding degradative systems (Broda et al., 2000a), bacteriocin production (Jones et al., 2009), and resistance to antibiotics (Sebaihia et al., 2006). These traits may improve their competitiveness by enabling faster genome replication through gene dosing effects and a higher growth rate of the bacterial cell. It is possible that extrachromosomal elements serve as vehicles for the exchange of genomic information between different strains and species or potentially to other genera such as lactic acid bacteria (LAB), that may provide a competitive advantage within their specific microbial ecosystem.

Comparative genome and glycobiome analyses have identified considerable variation in the conservation of orthologous gene families and CAZy profiles both between and within the meat spoilage associated *Clostridium* species (Fig. 3). This suggests a degree of specialization within these bacteria, especially with the presence and abundance of a set of genes required for polysaccharide degradation such as: CE4 and CE6 acetyl xylan esterases (EC 3.1.1.72); CE9 N-acetylglucosamine 6-phosphate deacetylases (EC 3.5.1.25); and an assortment of multi-modal chitin or peptidoglycan cleaving enzymes consisting of GH18 (EC 3.2.1.14) with CBM50; and GH13 with either CBM48 glycogen, CBM41 α-glucans amylose, amylopectin, pullulan, and oligosaccharide, or CBM34 starch-binding modules. The abundance of GH and CE domain-containing CAZymes encoded within the *C. estertheticum* DSM 8809^T^ and *C. estertheticum subsp. laramiense* DSM 14864^T^ genomes, in particular their PL complements (PL1 and PL9 pectate lyases (EC 4.2.2.2), PL4 and PL11 rhamnogalacturonan endolyases (EC 4.2.2.23)), suggests that they are specialist pectin and also xylan degraders. In addition to the CAZymes predicted to metabolize complex insoluble polysaccharides such as xylan and pectin, a large repertoire of enzymes were predicted to be encoded intracellularly (GH8, GH28, GH39, GH51, GH67, GH105, GH115, and CE2 families), in both bacteria. These findings are consistent with the phenotypic description and characterization of *Clostridium algidixylanolyticum* sp. nov. with the type strain SPL73^T^ (DSM 12273^T^), a psychrotolerant, xylan-degrading, spore-forming *bacterium* isolated from vacuum-packed, temperature-abused raw lamb (Broda et al., 2000a). This suggests that certain *Clostridium* species associated with meat spoilage are well equipped to utilize a variety of complex oligo- and monosaccharides resulting from extracellular hydrolysis, as they are transported and metabolized within the cell as substrates for growth.

The two subspecies of *C. estertheticum* currently recognized have previously been grouped into lactate producing and lactate non-producing strains (Spring et al., 2003; Yang et al., 2010). The diversity in fermentation products observed within each *C. estertheticum* subspecies suggests differences in the metabolic pathways when grown on a range of substrates (Fig. 4). The metabolic pathway characterization analysis imply that certain *Clostridium* species may have the ability to switch substrate utilization from a simple monosaccharide substrate like glucose to a complex polysaccharide such as glycogen when grown in meat juice medium.

To explain the potential roles of *C. estertheticum* subsp. *laramiense* DSM 14864^T^ and *C. estertheticum* DSM 8809^T^ as specialized pectin fermenters due to their possession of PL CAZy family enzymes, we investigated the genes involved in uronic acid metabolism. In this pathway, broken-down pectin or uronic acid components on xylans are released in the form of galacturonates and glucuronates, which are metabolized via 2-keto-3-deoxygluconate (KDG) rather than *via* the EMP pathway. The KDG is then converted to 2-keto-3-deoxygluconate phosphate (KDGP) by 2-dehydro-3-deoxygluconokinase and is then converted to pyruvate and glyceraldehyde-3-phosphate (GAP) by 2-keto-3-deoxygluconate 6-phosphate aldolase (Fig. 4). Both *C. estertheticum* genomes encode the enzymes required to convert both glucuronate and galacturonate through to pyruvate and potentially result in ATP production. Also in the case of *C. estertheticum* subsp. *laramiense* DSM 14864^T^ and *C. estertheticum* DSM 8809^T^, their CAZy profiles and in particular their PL and CE content, may also account for the offensive odours and production of gas commonly associated with these species. Our comparative genomics findings provide further evidence for the need to include genome sequencing as a prerequisite for the description of new *Clostridium* species. For future work, further phenotypic characterization and biochemical investigation to differentiate the metabolic activity in vacuum-packaged meat spoilage associated *Clostridium* is warranted.

## 5.0 Conclusions

The genome sequences of the *Clostridium* species reported here is a valuable resource for future studies investigating the bacterial genetic mechanisms associated with BPS. In order to improve the phylogenetic resolution of the *Clostridium* genera and improve our limited knowledge of meat spoilage caused by *Clostridium* species, future efforts should focus on the generation of complete genomes across a wider range of *Clostridia* species.

## Acknowledgements

The research outlined in this study was supported by the AgResearch Ltd Strategic Science Investment Fund (SSIF), contract A25980.

